## Supplementary figures and images for "Molecular mapping and functional validation of GLP-1R cholesterol binding sites in pancreatic beta cells"

### Supplementary Figure 1

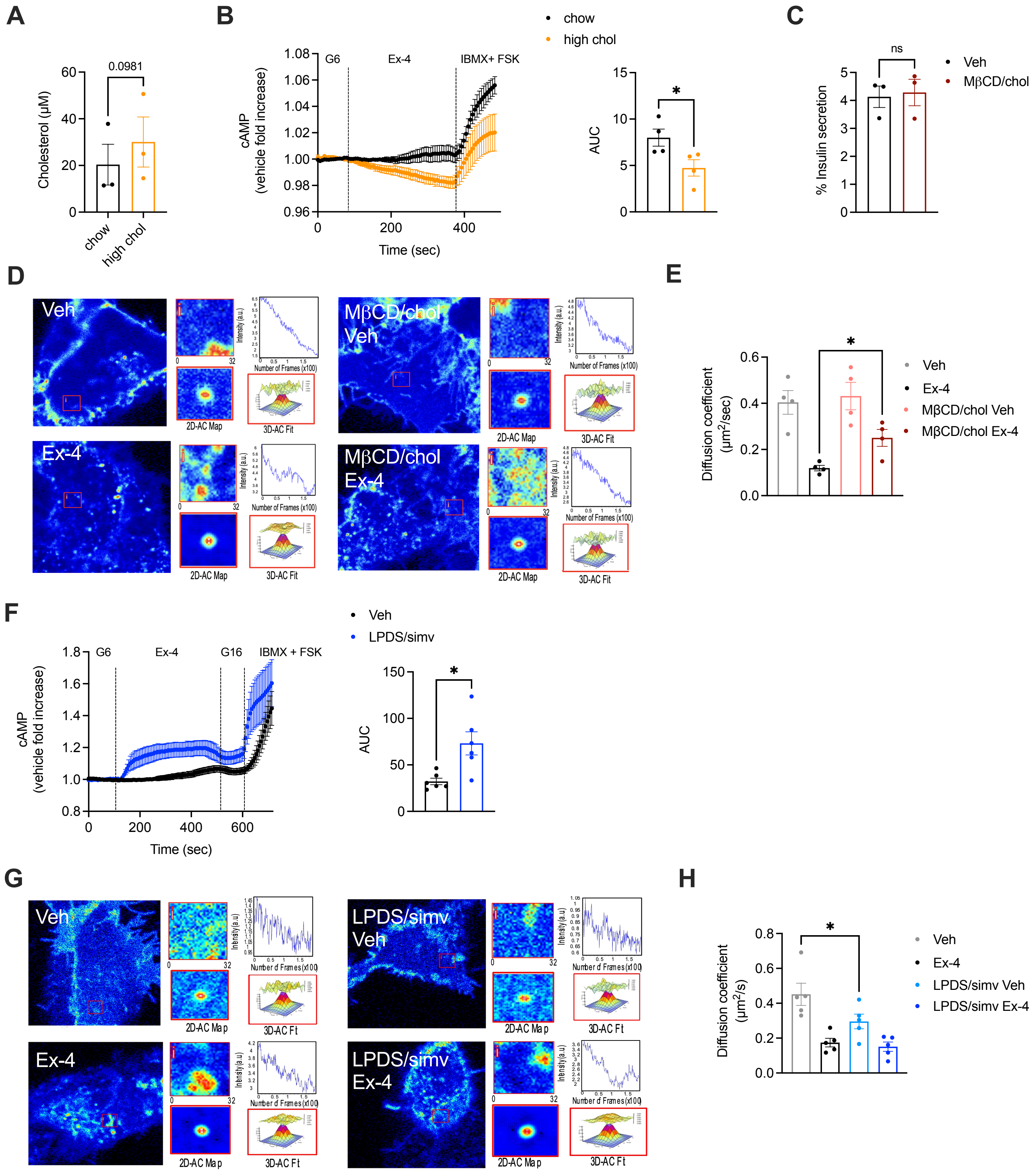

### Supplementary Figure 2

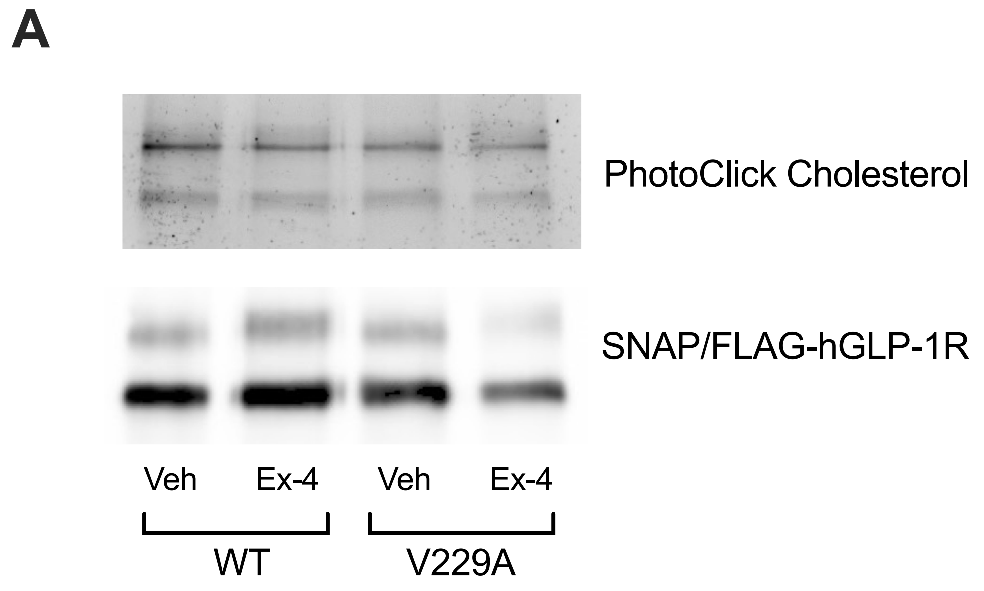


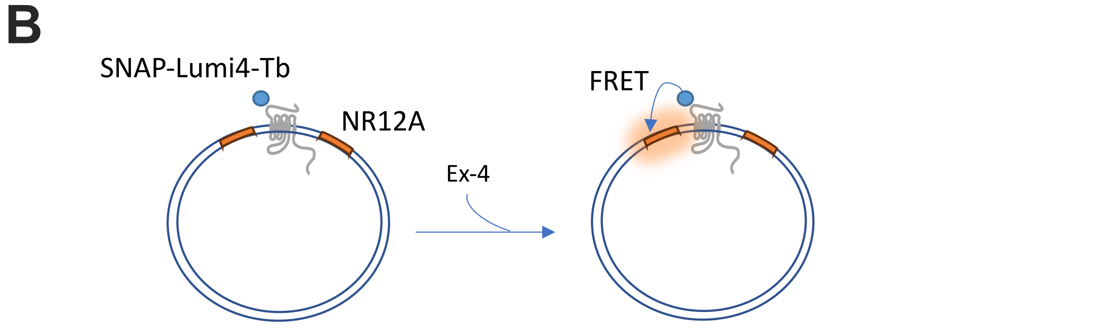


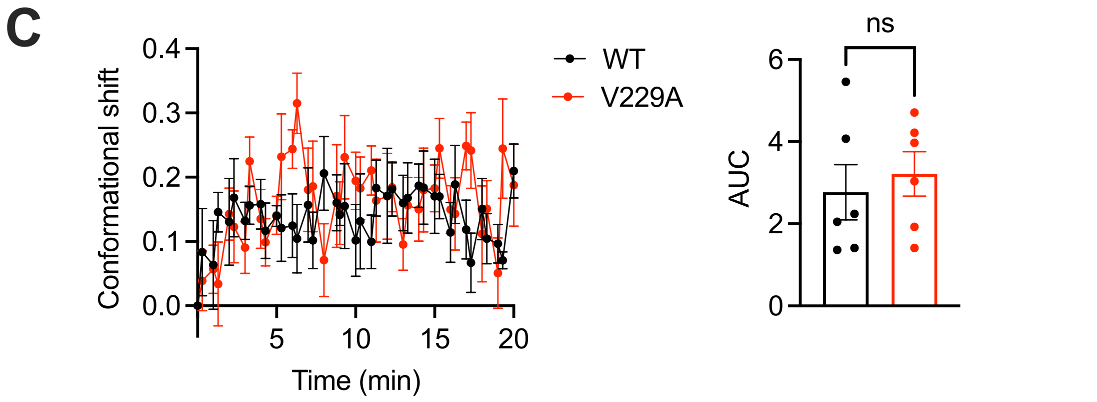


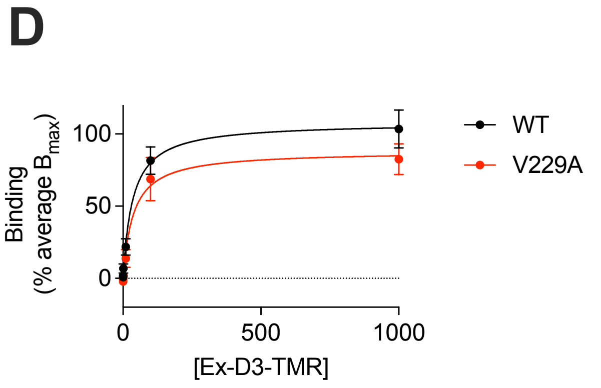


| | WT | V229A |
|---|---|---|
| Kd | 41 ± 10 | 55 ± 20 |

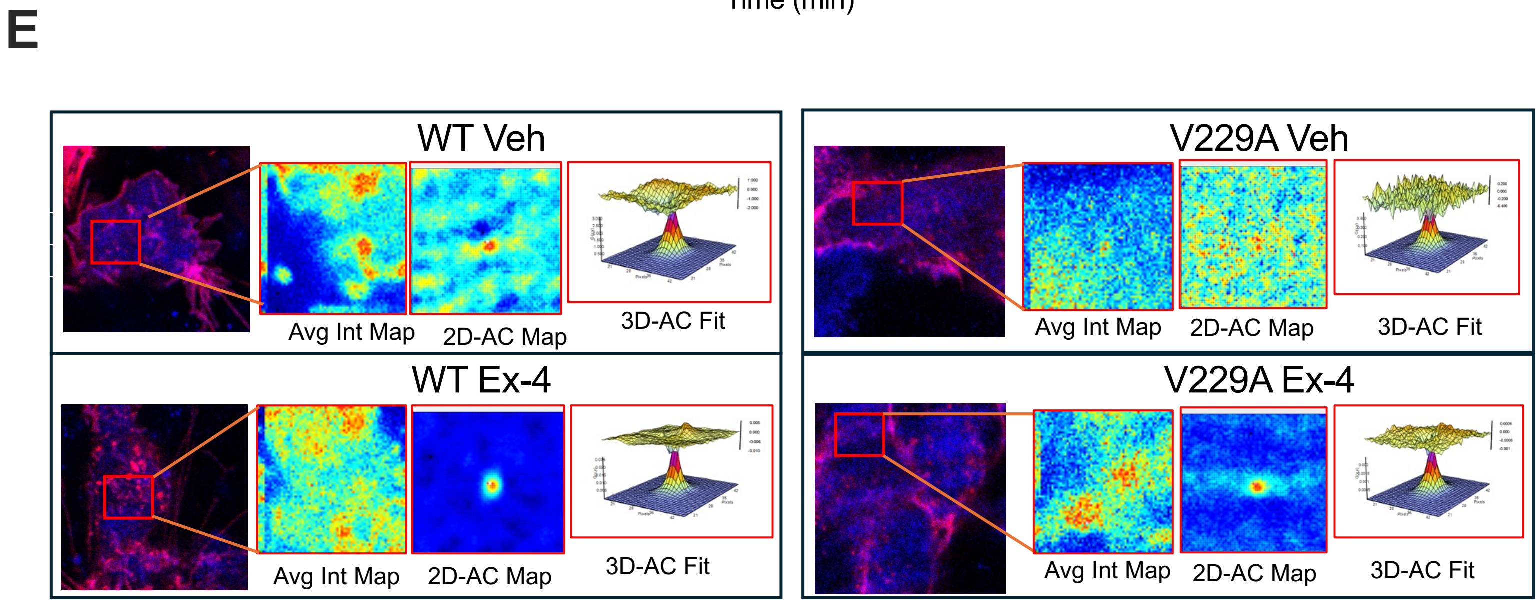

### Supplementary Figure 3

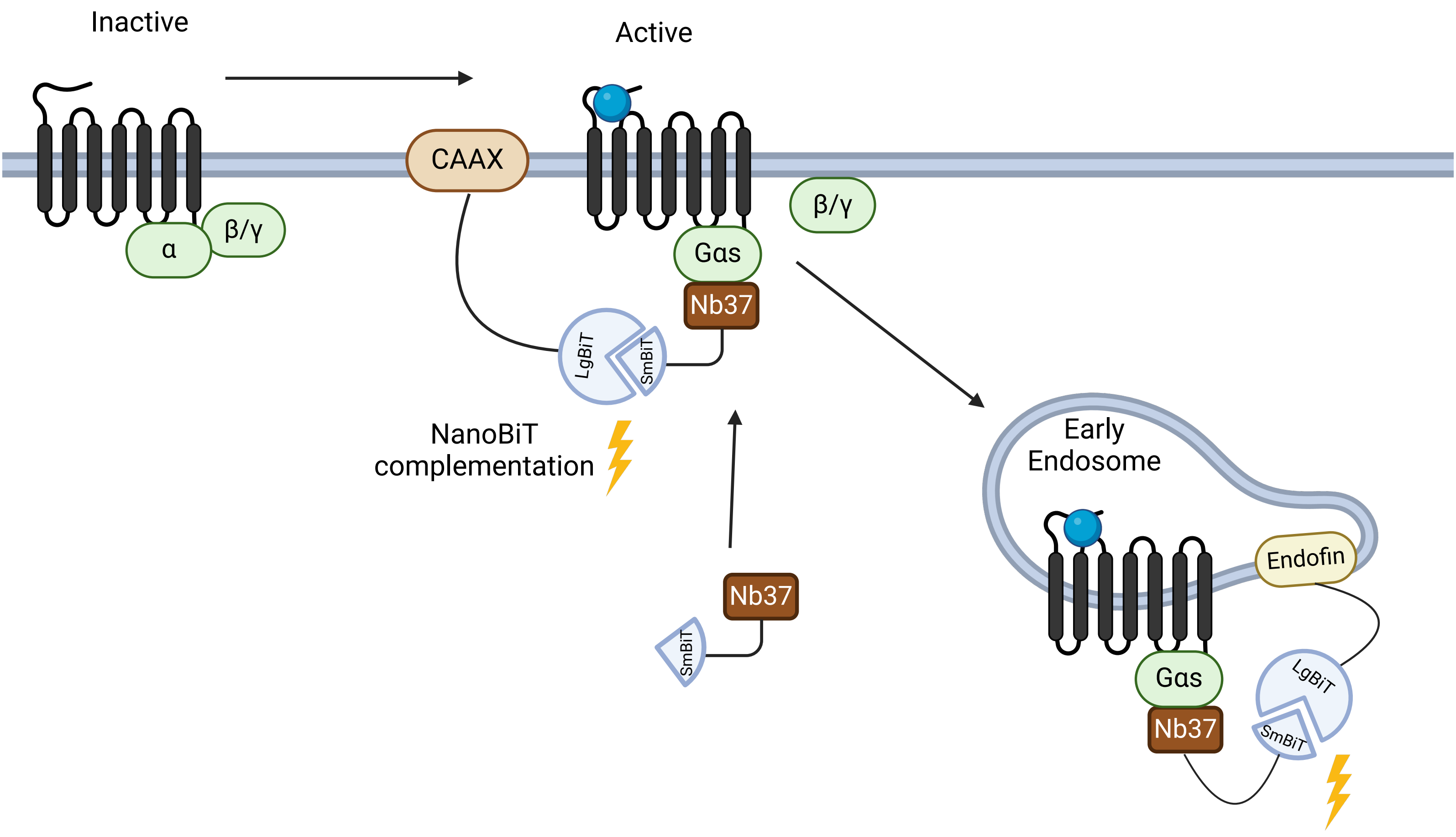

### Supplementary Figure 4

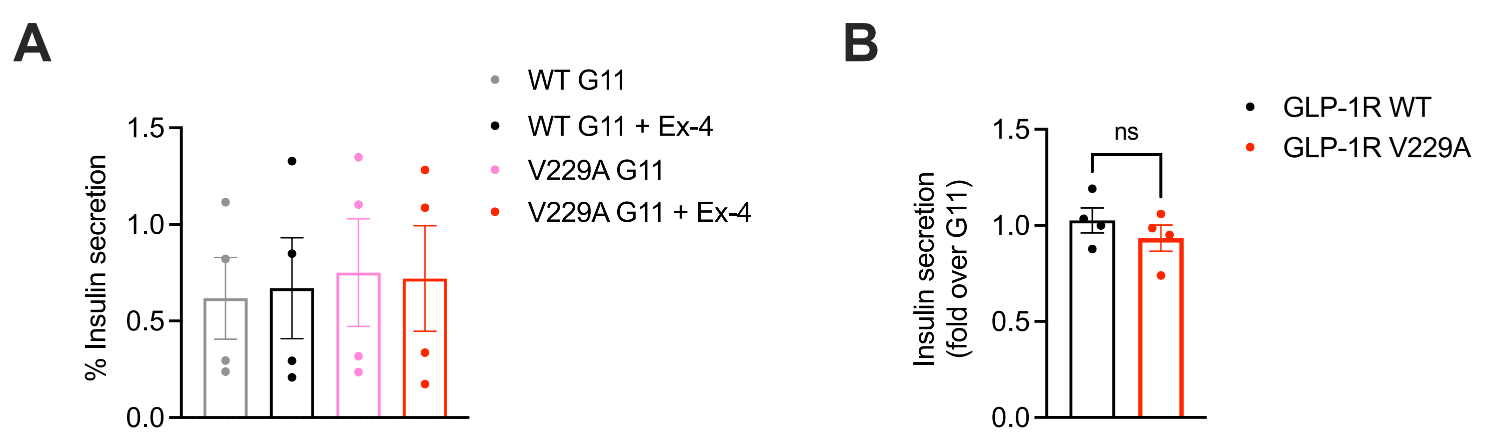
